# Genetic Signatures of Lipid Metabolism Evolution in Cetacea

**DOI:** 10.1101/250597

**Authors:** Yoshinori Endo, Ken-ichiro Kamei, Miho Inoue-Murayama

## Abstract

In mammalian evolutionary history, Cetacea (whales, dolphins, and porpoises) achieved astonishing success by adapting to an aquatic environment. One unique characteristic of cetaceans, contributing to this adaptive success, is efficient lipid utilization. Here we report comparative genetic analysis of aquatic and terrestrial Cetartiodactyla using 144 genes associated with lipid metabolism. We analyzed mutation rate, amino acid substitution, and metabolic pathways using genetic data publicly available. Our test detected 18 positively selected genes in Cetacea compared to 13 in Bovidae with little overlap between the lineages. We identified lineage-specific patterns of amino acid substitutions and functional domain that were mutually exclusive between cetaceans and bovids, supporting divergent evolution of lipid metabolism since the divergence of these taxa from a common ancestor. Moreover, a pathway analysis showed that the identified genes in cetaceans were associated with lipid digestion, lipid storage, and energy producing pathways. This study emphasizes the evolutionary context of lipid metabolism modification of cetaceans and provides a foundation for future studies of elucidating the adapted biological mechanisms of cetacean lipid metabolism and a framework for incorporating ecological context into studies aimed at investigating adaptive evolution.

## Introduction

The evolution of cetaceans represents one of the most striking adaptations for a habitat transition in mammalian evolutionary history (Thewissen et al. 2009; Uhen 2010). Although mammals are generally under greater constraints with regard to anatomical, physiological, and behavioral changes than other classes of vertebrates, cetaceans have achieved a remarkable macroevolutionary transition since their divergence from an artiodactyl ancestor approximately 50 million years ago (Thewissen et al. 2009). The past two decades of paleontological and phylogenetic research have thoroughly characterized the evolutionary history of cetaceans (Thewissen et al. 2009; Uhen 2010; Gatesy et al. 2013), and there continues to be considerable attention on the molecular evolution behind the dramatic changes in their morphology and ecology upon aquatic adaptation (McGowen et al. 2014). Understanding the molecular basis underlying the adaptive traits will shed light on this unique evolutionary event from land to water.

Cetaceans have developed unique characteristics that are different from those of terrestrial mammals but are common within the lineage (Thewissen 2014). For example, all species of Cetacea are carnivores and have streamline body morphology. Cetaceans consumes lipids not only as one of their primary energy resources, but also store lipids as an important component of the thick blubber that provides thermo insulation and supports their streamlined morphology (Berta et al. 2006). The evolution of lipid usage may be a key factor for the adaptive characteristics enabling cetacean transition to an aquatic environment. Recent genomic studies have reported signatures of positive selection on diverse processes, including those of lipid metabolism (McGowen et al. 2012; Sun et al. 2013; Foote et al. 2015; Tsagkogeorga et al. 2015; Wang et al. 2015). However, evolutionary analyses of cetaceans have been with distantly related lineages, rather than contrasting them with closely related terrestrial species using the same analytical procedures. Furthermore, given the potential adaptive specificity of each cetacean species to its own environment and ecology, analyses at the taxonomic-group level are necessary to interpret the detected genetic signatures as aquatic adaptations.

In this study, we identified the genetic signatures of positive selection on lipid metabolism in Cetacea and Bovidae (Fig. 1*a*). The goals of our study were to (1) compile genes associated with lipid metabolism from Cetartiodactyla species (even-toed ungulates, including Cetacea), (2) identify the positively selected genes (PSGs) for each lineage, Cetacea and Bovidae, (3) determine the amino acid substitutions that occurred in functional domain regions of the identified PSG, and (4) describe the biological functions and metabolic pathways that have been modified during aquatic adaptation (Fig. 1*b*). Our comparative analyses revealed novel and strong signatures for positive selection in the cetacean lipid metabolism pathways compared with those of bovids. This study provides the genetic basis for the evolution of lipid metabolism developed by cetaceans in the context of aquatic adaptation and emphasizes the importance of comparative analyses based on the phylogenetic and ecological perspectives of the study organisms.

**Figure 1.**
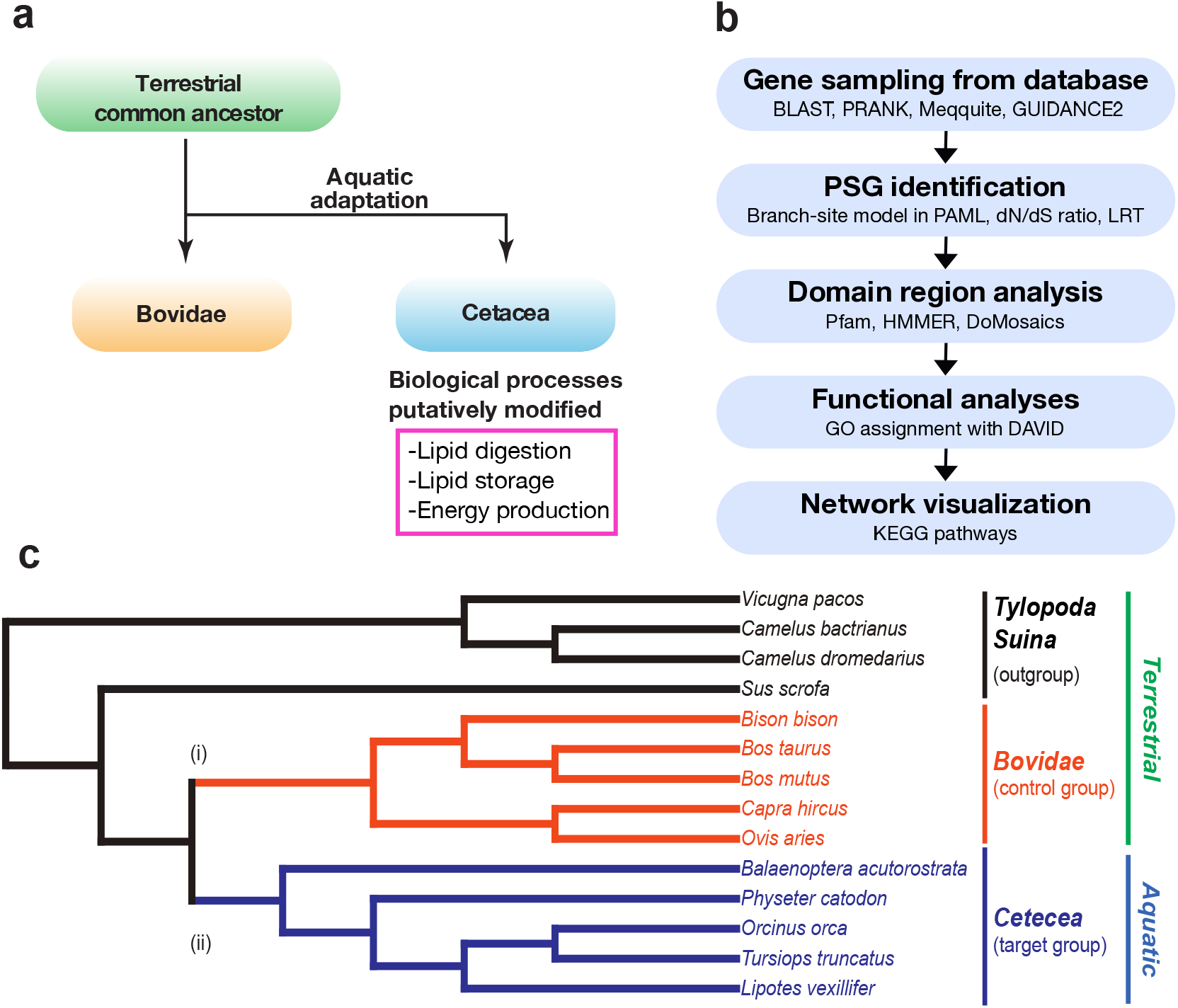
Genetic signatures of aquatic adaptation in an evolutionary context. (a) Diagram describing the evolutionary relationship between Cetacea and Bovidae. Cetacea and Bovidae diverged from a terrestrial ancestor and adapted to aquatic and terrestrial habitats. The adaptive features for the aquatic environment indicate rearrangement of biological processes associated with lipid metabolism. (b) Schematic illustration of methods used to investigate the adaptive signatures of Cetacea lipid metabolism. (c) Cladogram showing the evolutionary relationships among Cetartiodactyla species. Branch color represents the lineages: five Cetacea (*blue*: target aquatic group), five Bovidae (*orange:* control terrestrial group), and one Suina and three Tylopoda (*black:* outgroup). The phylogenetic positions of the species were reconstructed according to previous works (Agnarsson and May-Collado 2008; Gatesy et al. 2013). The lineages tested for positively selected genes are indicated with (i) and (ii).

## Materials and Methods

### Data sampling and preparation

All the genetic data in this study were downloaded from NCBI (https://www.ncbi.nlm.nih.gov). Genomes of Cetartiodactyla species that were publicly available and annotated were used. Orthologous cording DNA sequences (CDSs) were identified by blastn search with human genes as references. Genes that were not annotated in any of the study species were excluded from later analyses. CDSs were aligned with PRANK v. 140603 (Löytynoja and Goldman 2005) and manually edited Mesquite v.3.2 (Maddison and Maddison 2017). To remove potentially unreliable sequences, the aligned CDSs were filtered with GUIDANCE2 using the default settings (Sela et al. 2015). In addition, codon triplets with gaps in more than 50% of the were discarded.

### Identification of positively selected genes

PSGs were identified using branch-site models implemented in the CODEML program the PAML v. 4.8 software package (Yang 2007). The branch-site test detects positive selection on *a priori* specified branches. To detect selection in Cetacea and Bovidae separately, we conducted the tests twice for all genes evaluating one lineage at a time. A one-ratio model was employed as a null hypothesis in which all branches were under neutral evolution. A two-ratio model was conducted as an alternative hypothesis in a specified lineage was under positive selection, and this model estimated the *d_N_/d_S_* ratio (ω: nonsynonymous to synonymous substitution ratio). Because the branch-site test estimates the *d_N_/d_S_* ratio on a subset of codons, the ω values are not reliable measure of the strength of positive selection (Yang and dos Reis 2011). Therefore, the likelihood ratio test was used to determine whether a gene was under positive selection. The likelihood ratio test was performed to compare 2Δ*l* of the two models to the χ^2^ distribution for *p*-value evaluation.

### Domain annotation and amino acid substitutions

The identified PSGs were scanned for domains with the pfam_scan utility and HMMER3.1 (Finn et al. 2015) against the Pfam-A from Pfam v. 31.0 (Finn et al. 2016). The domain arrangements were visualized with DoMosaics v. 0.95 (Moore et al. 2014). Domain similarity was estimated by using amino acid sequences of *Orcinus orca* and *Bos taurus* as criteria for Cetacea and Bovidae PSGs, respectively, using DoMosaics, and the dot plots were generated using the R package ggplot2 (Wickham 2009). The lineage-specific amino acid substitutions were identified manually by looking at a conversion of residue that occurred in only one lineage and where all the species in that lineage had identical residue. Those instances were included when one of the outgroup species had a mutation at the same position.

### Function and pathway analysis of PSGs

Gene ontology (GO) terms of the identified PSGs were assigned according to DAVID Functional Annotation tool (Huang et al. 2009b, a) under the biological process domain. The pathways in which the identified PSGs participate were identified using KEGG Mapper (Kanehisa et al. 2017). Because a gene can be involved in multiple pathways, which can be categorized in multiple hierarchies, the pathways that was more representative and directly related with lipid metabolism were focused.

## Results and Discussion

### Positively selected genes in Cetacea and Bovidae

To detect signatures of positive selection, we analyzed the alignments of lipid metabolism-related genes collected from genomes of Cetartiodactyla. To assess the evolutionary transitions since their divergence from a terrestrial ancestor (Thewissen et al. 2009), we categorized the study species by their phylogeny and ecology in the target group (Cetacea), a control group (Bovidae), and an outgroup (Suina and Tylopoda). Among Ruminantia, the family Bovidae was used as a control group. Bovidae is a fully terrestrial and closely related clade to Cetacea (Shimamura et al. 1999). The ecological and geographical diversity of bovids provides an unbiased representation of terrestrial adaptation. We opted to use only one family, because there are few differences in the adaptive fitness of bovids to an aquatic environment. Thus, positive selection detected in Cetacea can suggest the adaptations required to return to an aquatic environment. To gain insight into the evolution of lipid usage, we investigated the genes associated with lipid metabolism. Taken together, the results from these datasets will elucidate plausible molecular adaptations underlying cetacean macroevolutionary transitions from a terrestrial to an aquatic environment.

A total of 14 Cetartiodactyla species were used in this study: five Cetacea, including the minke whale (*Balaenoptera acutorostrata*), sperm whale (*Physeter catodon*), killer whale (*Orcinus orca*), bottlenose dolphin (*Tursiops truncatus*), and Yangtze River dolphin (*Lipotes vexillifer);* five Bovidae, including cattle (*Bos taurus*), wild yak (*Bos mutus*), American bison (*Bison bison*), domestic goat (*Capra hircus*), and domestic sheep (*Ovis aries);* and Suina and Tylopoda, including pig (*Sus scrofa*), alpaca (*Vicugna pacos*), Bactrian camel (*Camelus bactrianus*), and Arabian camel (*Camelus dromedaries*) (Fig. 1c). For each species we searched orthologous of the 156 genes assigned to functions associated with lipid metabolism in humans. Our filtering strategy retained 144 genes covering the major lipid metabolic pathways (Table S1).

To identify genes under positive selection, we performed likelihood ratio tests using branch-site models implemented in PAML (Yang 2007). This analysis estimates *d_N_/d_S_* ratio (ω) for the lineage of interest, and the PSGs were determined by the likelihood ratio test. Of the 144 lipid metabolism genes, 14 genes were identified as PSG in the lineage of Cetacea, as well as 5 genes in the lineage of Bovidae at *p* < 0.05 (Fig. 2*a*, Table 1, and also Table S2 and S3). Due to the low number of PSGs for assessing the selection landscape of metabolic pathways, we relaxed the significance level to 0.05 ≤ *p* < 0.1 and found an additional 4 and 8 PSGs in Cetacea and Bovidae, respectively. Among the identified PSGs, only one gene, *GPAM* (also known as *GPAT1*, glycerol-3-phosphate acyltransferase) was found in common at *p* < 0.05 (Fig. 2*b*).

**Table 1.** List of positively selected genes (PSGs) identified in Cetacea and Bovidae.

| Lineage | Gene symbol | Estimates of $\omega$ | $\kappa$ | lnL one-ratio | lnL two-ratio | p value |
| --- | --- | --- | --- | --- | --- | --- |
| Cetacean | *CYP8B1* | 11.92734 | 5.85792 | -4291.819841 | -4285.67461 | 4.55.E-04 ** |
|  | *PDHB* | 5.98748 | 4.36259 | -2494.18574 | -2491.079352 | 1.27.E-02 * |
|  | *AGPAT1* | 7.05413 | 4.86528 | -1855.189217 | -1852.248895 | 1.53.E-02 * |
|  | *LIPF* | 4.44878 | 2.25004 | -6287.045085 | -6284.171286 | 1.65.E-02 * |
|  | *PNLIPRP2* | 4.54093 | 3.64754 | -4524.349375 | -4521.629714 | 1.97.E-02 * |
|  | *APOB* | 4.05216 | 4.19478 | -41746.17128 | -41743.46198 | 1.99.E-02 * |
|  | *CYP2J2* | 13.63547 | 3.19121 | -5139.202241 | -5136.697766 | 2.52.E-02 * |
|  | *EHHADH* | 14.52325 | 3.44028 | -5810.991244 | -5808.631236 | 2.98.E-02 * |
|  | *ACADVL* | 3.40418 | 4.11549 | -5098.363465 | -5096.023469 | 3.05.E-02 * |
|  | *GPAM* | 4.35956 | 4.01882 | -6046.804276 | -6044.470464 | 3.07.E-02 * |
|  | *CYP2E1* | 2.37475 | 4.51029 | -4844.274535 | -4842.07278 | 3.59.E-02 * |
|  | *HADHB* | 6.58348 | 4.73191 | -3491.684049 | -3489.483724 | 3.59.E-02 * |
|  | *APOA2* | 3.60699 | 4.43456 | -961.661901 | -959.657031 | 4.52.E-02 * |
|  | *PGD* | 3.30916 | 3.05453 | -3813.94106 | -3812.008851 | 4.93.E-02 * |
|  | *GNPAT* | 2.10957 | 4.04115 | -5941.958899 | -5940.047374 | 5.06.E-02 |
|  | *ACSS2* | 8.09817 | 4.63572 | -4902.585196 | -4900.799437 | 5.88.E-02 |
|  | *PNLIP* | 3.79609 | 3.17642 | -4448.612671 | -4446.922816 | 6.60.E-02 |
|  | *FBP1* | 2.87282 | 3.22927 | -2728.680035 | -2727.07769 | 7.34.E-02 |
| Bovidae | *MOGAT3* | 6.44231 | 3.04369 | -4562.007537 | -4552.517873 | 1.32.E-05 ** |
|  | *ABHD12* | 10.698 | 4.10908 | -2503.418086 | -2499.826239 | 7.36.E-03 ** |
|  | *APOE* | 32.57059 | 4.04548 | -3013.622556 | -3010.088442 | 7.85.E-03 ** |
|  | *MLYCD* | 12.46396 | 4.31351 | -4458.618387 | -4456.297488 | 3.12.E-02 * |
|  | *GPAM* | 8.83467 | 4.03293 | -6050.187322 | -6047.92294 | 3.33.E-02 * |
|  | *IDH2* | 45.30329 | 6.95175 | -3056.751425 | -3054.899266 | 5.43.E-02 |
|  | *HSD17B4* | 16.02856 | 4.15483 | -5827.966765 | -5826.198171 | 6.00.E-02 |
|  | *SOAT1* | 45.93865 | 3.48586 | -4577.415397 | -4575.845606 | 7.64.E-02 |
|  | *PGAM5* | 22.60281 | 5.77771 | -1980.355128 | -1978.812842 | 7.90.E-02 |
|  | *ACAD9* | 2.64777 | 4.91697 | -5713.112028 | -5711.57532 | 7.96.E-02 |
|  | *MOGAT1* | 5.222 | 3.44578 | -2989.32785 | -2987.877786 | 8.86.E-02 |
|  | *ACADM* | 21.5195 | 2.95373 | -2911.486719 | -2910.067882 | 9.21.E-02 |
|  | *SOAT2* | 12.78966 | 4.68574 | -4299.927829 | -4298.536813 | 9.53.E-02 |

Notes.— $\kappa$ , transition/transversion rate ratio; ln L one-ratio and ln L two-ratio, log likelihood value for null and alternative hypothesis, respectively; \*, significant ( $P < 0.05$ ); \*\*, highly significant ( $P < 0.01$ ).

**Figure 2.**
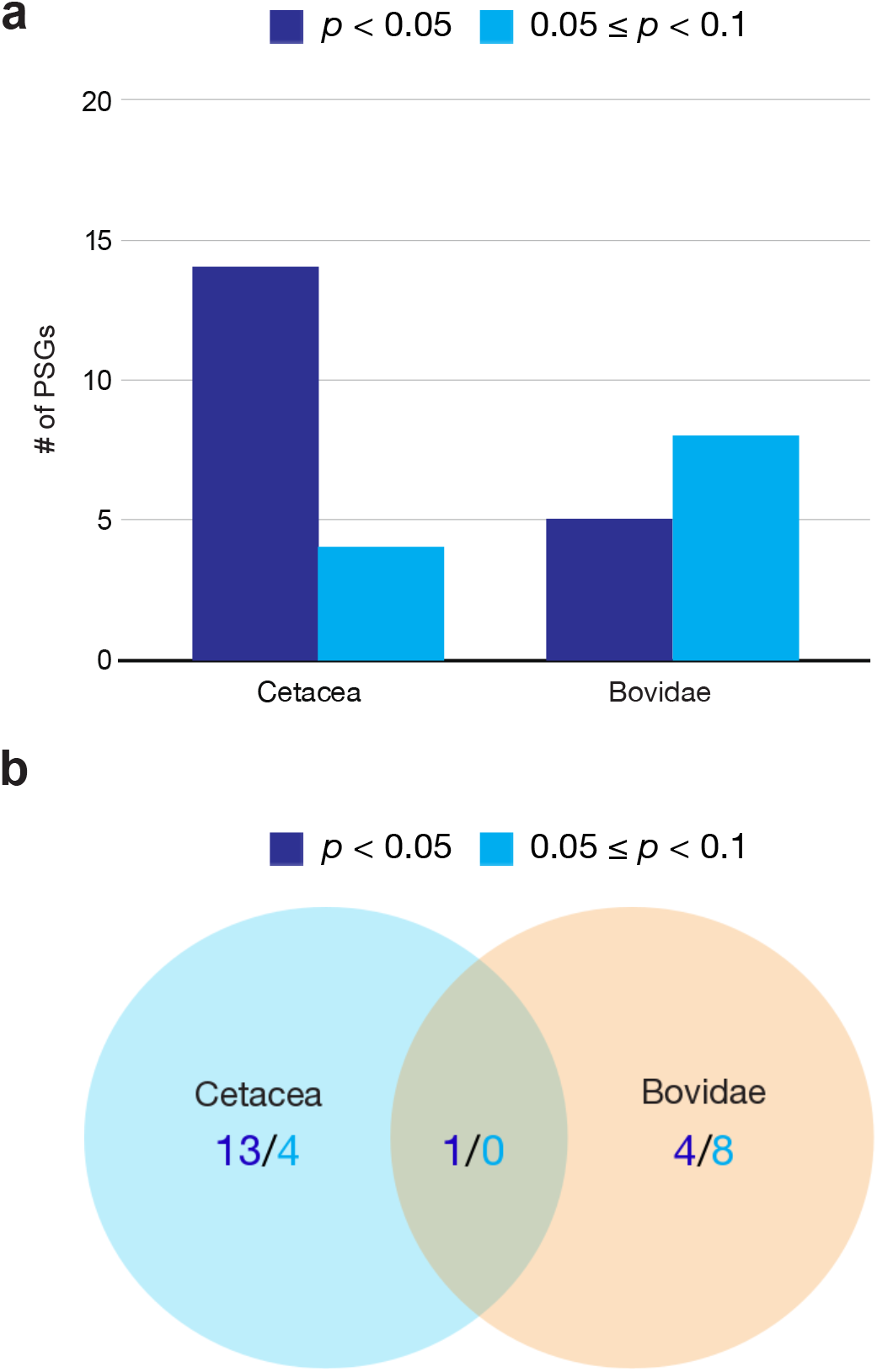
Number of identified positively selected genes (PSGs). (a) Bar graph showing the number of PSGs in Cetacea and Bovidae (*blue*: *p* < 0.05; *light blue:* 0.05 ≤ *p* < 0.1). (b) Venn diagram showing unique and overlapping PSGs in Cetacea and Bovidae (*blue*: *p* < 0.05; *light blue:* 0.05 ≤ *p* < 0.1).

To assess whether the detected amino acid substitutions have influence on functions of the identified PSGs, we examined the functional regions of the protein sequences, namely domains, as described in the Method section (Fig. 3 for *EHHADH*; for other PSGs see Fig. S1-S31). Whereas the domain arrangements were largely conserved among Cetartiodactyla, the comparisons of the sequences of the domain region showed inter-lineage divergence between Cetacea and Bovidae. Furthermore, we explored amino acid substitutions that occurred only in Cetacea and remain identical within the lineage. Such lineage-specific substitutions in the functional domain of genes, evolving under positive selection, would putatively be associated with adaptive phenotype in an aquatic environment. We found lineage-specific substitutions in at least one domain of 14 Cetacea PSGs and 11 Bovidae PSGs (Table 2 and Fig. 3*c*). For example, one Cetacea PSG, *EHHADH*, is composed of four domains, except for the domain arrangements of *Tursiops truncates* and *Sus scrofa*, probably because of isoforms (Fig. 3*a*). The amino acid sequence of each domain differs between Cetacea and Bovidae while they are relatively similar within each lineage (Fig. 4*b*). *EHHADH* encodes two enzymes, enoyl-CoA hydratase/isomerase (ECH) and 3-hydroxyacyl-CoA dehydrogenase (HCDH), involved in β-oxidation (Bartlett and Eaton 2004). The amino acid sequences of both of these enzymes contains lineage-specific substitutions (Fig. 4*c*), even though none of the substitutions were found in active sites (Agnihotri and Liu 2003). We did not find lineage-specific substitutions in four Cetacea PSGs (*AGPAT, FBP1, GNPAT*, and *GPAM*) nor in two Bovidae PSGs (*GPAM* and *PGAM5*). However, substitutions outside of domains may still have functional influence.

**Table 2.**
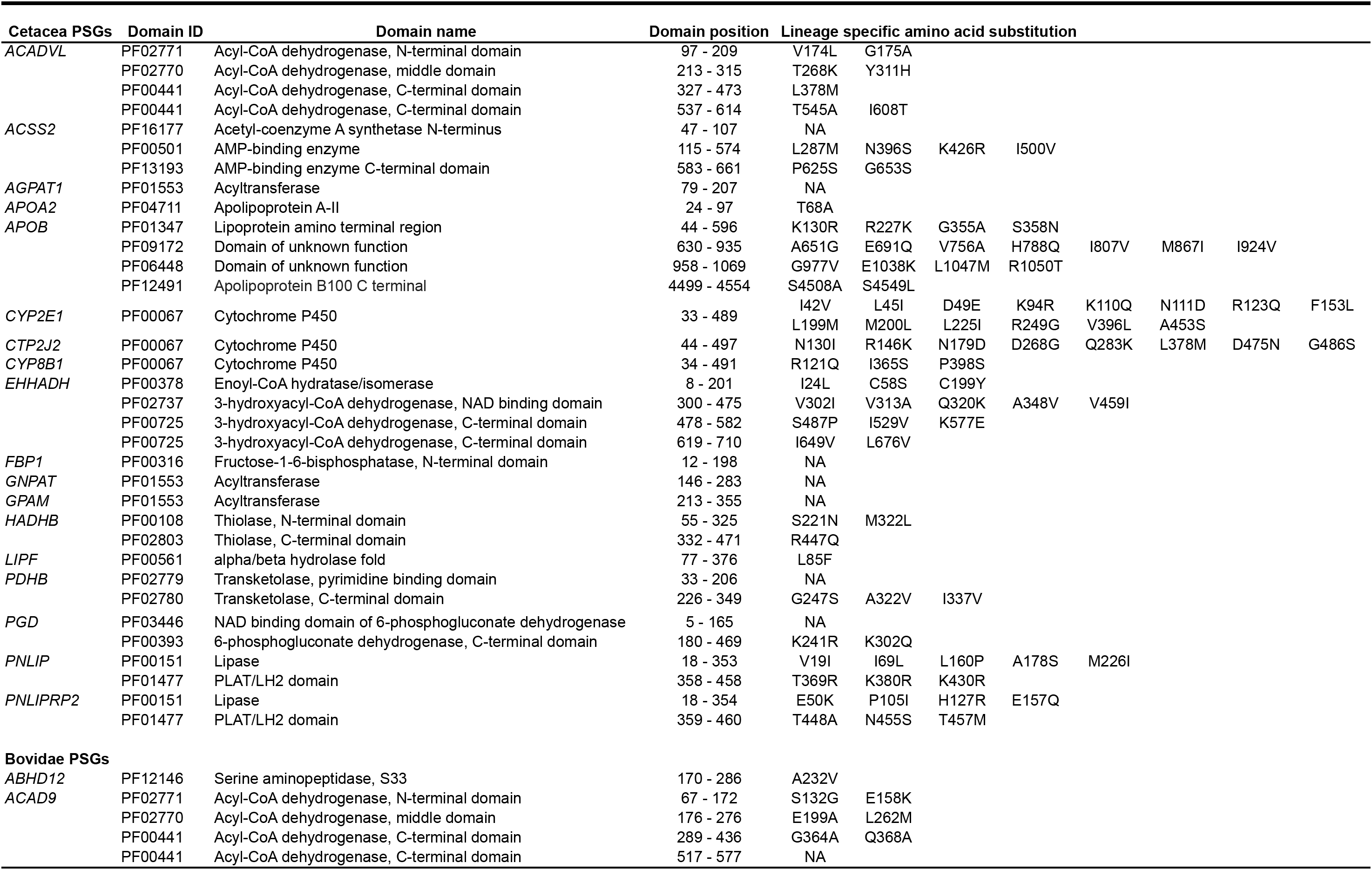
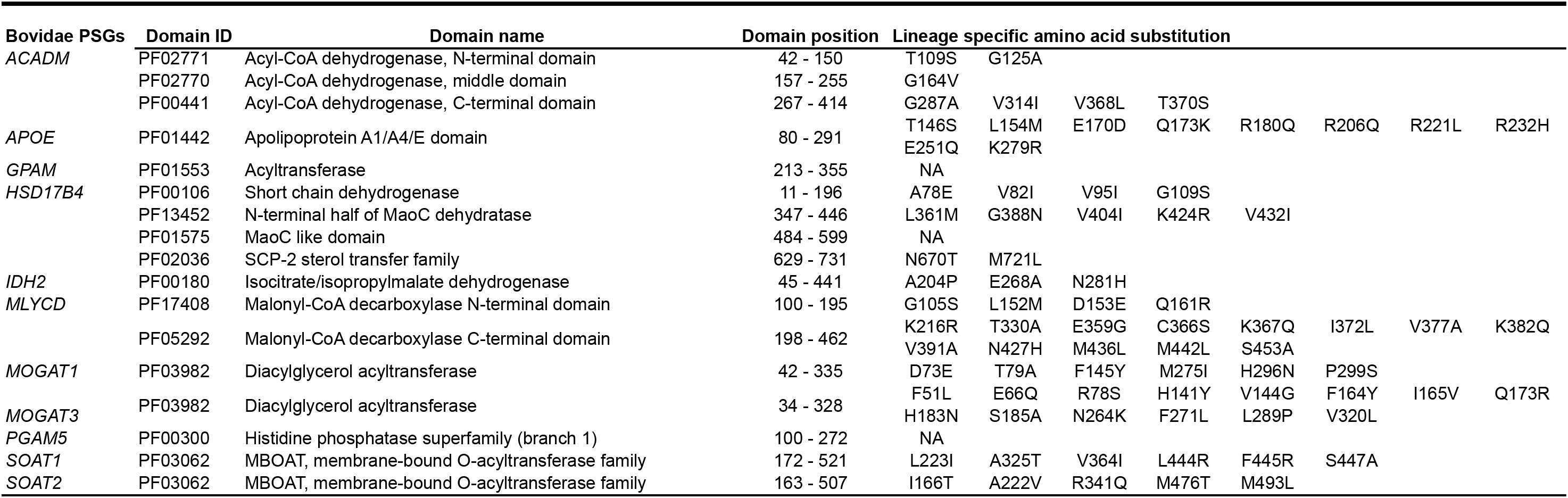
Lineage specific amino acid substitutions that occurred in domain regions of positively selected genes (PSGs)

**Figure 3.**
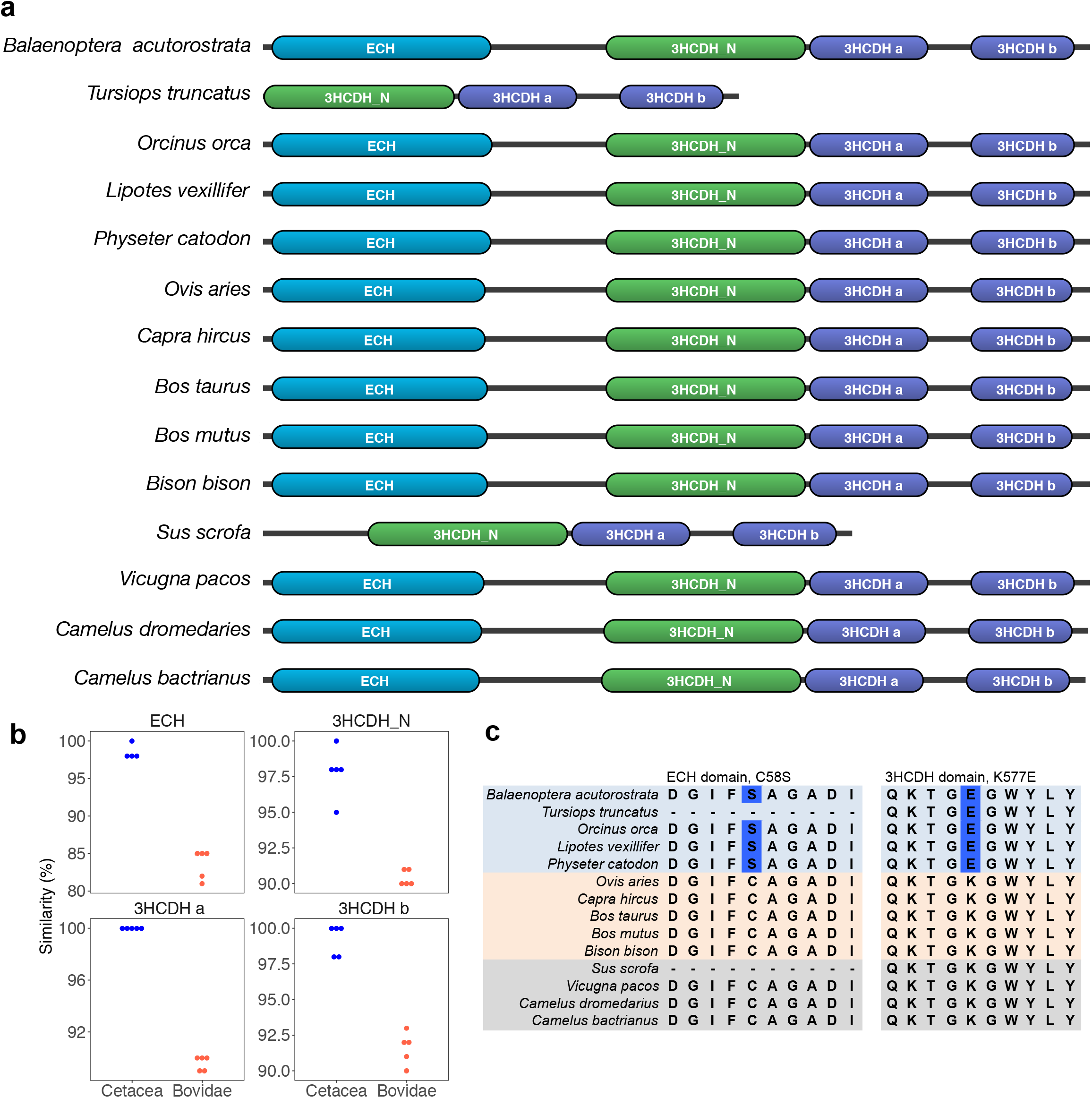
Amino acid substitutions that were found in functional domains of positively selected genes (PSGs), *EHHADH* as an example. (a) Domain arrangements showing the name and position of domains. (b) Dot plots for the amino acid sequence similarity of each domain. *Blue* dots represent cetacea species, and *orange* dots represent bovidae species. (c) Lineage specific amino acid substitution that were unique to and identical within Cetacea.

Our comparison of the lipid metabolism genes highlighted greater and unique selective pressures in cetacean lipid metabolism compared with Bovidae. The results showed that both lineages have modified their lipid metabolism since divergence from a common ancestor, suggesting the adaptive importance of the usage of lipids in both aquatic and terrestrial environments. The relatively higher number of PSGs with *p* < 0.05 within Cetacea, however, supports the hypothesis that evolutionary changes in lipid metabolism were more substantial for the aquatic than the terrestrial lineage. Moreover, the low extent of overlap between the Cetacea and Bovidae PSGs indicated the differing selection regimes in lipid metabolism of aquatic and terrestrial lineages. Our results (1) provide support for the adaptive evolution of lipid metabolism in the context of aquatic adaptation and (2) highlighted that these adaptations are not species specific for *T. truncatus* (Nery et al. 2013; Sun et al. 2013) or other cetacean species (Wang et al. 2015).

### Functional analysis of Cetacea PSGs

To gain insights into the functional consequences of the lipid metabolism behind the aquatic adaptation of cetaceans, we assessed the GO terms and metabolic pathways of the identified PSGs (Fig. 4).

**Figure 4.**
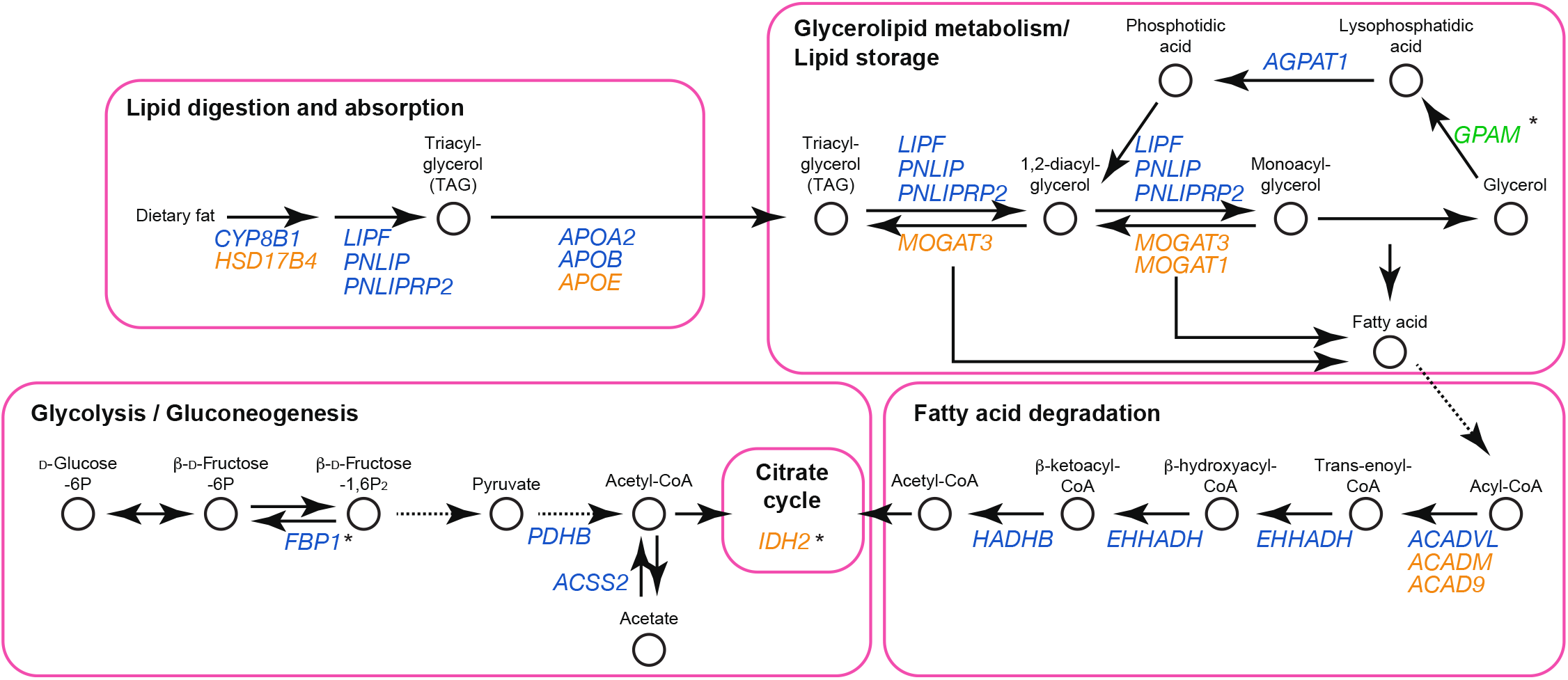
Unique and overlapping metabolic pathways based on gene ontological (GO) terms of the positively selected genes (PSGs) in Cetacea (*blue*), Bovidae (*orange*), and both (*green*). Solid arrow represents transformation of the substrates in one-step, and dashed arrow represents multiple steps. Each asterisk represents the rate limiting step. Each metabolic pathway is indicated in a *pink* rounded-square.

Our functional analyses revealed that six Cetacea PSGs (*APOA2, APOB, CYP8B1, LIPF, PNLIP*, and *PNLIPRP2*) were involved in the fat digestion and absorption pathway associated with the lipid catabolic process (GO:0016042) and lipoprotein biosynthetic process (GO: 0042158). *LIPF, PNLIP*, and *PNLIPRP2* encode pancreatic lipase, which hydrolyses dietary lipids in the digestive system (Pafumi et al. 2002; Mukherjee 2003). *CYP8B1* encodes a member of the cytochrome P450 superfamily of enzymes that catalyzes the synthesis of primary bile acids. Bile acids play important roles not only in absorption of lipids but also in the regulation of lipid metabolism. Additionally, they are believed to function as a therapeutic approach for metabolic syndrome (Ma and Patti 2014). Lastly, *APOA2* and *APOB* encode apolipoproteins, which aid in transporting hydrophobic lipids through the circulation system and are associated with atherosclerotic cardiovascular diseases (Grundy 2016), implying enhanced tolerance for high lipid concentration in their circulatory systems. Previous studies have also reported some genes in this pathway under selection in cetaceans (Sun et al. 2013; Foote et al. 2015; Wang et al. 2015; Wang et al. 2016); moreover, owing to the explicit comparison with their herbivorous relatives, our results provide further support of genetic signatures in cetacean adaptation in their shift in diet.

The storage of fat is an essential part of lipid metabolism, whereby triacylglycerol (TAG) is synthesized to serve as a major component in several biological functions. Synthesis of TAG takes place in glycerophospholipid metabolism via two distinct major pathways, the phosphatidic acid pathway and the monoacylglycerol pathway (Takeuchi and Reue 2009). We found five Cetacea PSGs (*AGPAT1, GPAM, LIPF, PNLIP*, and *PNLIPRP2*) involved in storage of fat, including the triglyceride metabolic process (GO: 0006641) and phosphatidic acid biosynthetic process (GO: 0006654). Three Bovidae PSGs (*GPAM, MOGAT1*, and *MOGAT3*) were also found in this category, including the glycerol metabolic process (GO: 0006071), in which TAG is synthesized from monoacylglycerol. These findings suggest enhanced TAG syntheses in both lineages, but under different molecular mechanisms in aquatic and terrestrial Cetartiodactyla. Functional evolution of the monoacylglycerol pathway associated with rumen evolution, for digesting C4 grasses that emerged approximately 40 million years ago, has been proposed for Bovidae (Jiang et al. 2014). Conversely, an evolutionary analysis of cetaceans reported positive selection on genes involved in both pathways of TAG synthesis for blubber thickening (Wang et al. 2015). In these studies, one cannot rule out that changes in the monoacylglycerol pathway may have been present in a common ancestor (i.e. contemporaneous evolution) since close relatives were not investigated. Our analyses do not suffer from this shortcoming because both Cetacea and Bovidae were analyzed in an equivalent manner. Therefore, our analyses support the evolution of the monoacylglycerol pathway in Bovidae and the phosphatidic acid pathway in Cetacea. The only common PSG between Cetacea and Bovidae, *GPAM*, encodes a mitochondrial enzyme that regulates and synthesizes TAG and other lipids (Wendel et al. 2009). More research, such as *in vitro* functional experiments, is necessary to determine whether the mutations in this gene produced a different phenotypic outcome between the lineages.

We also inspected evolutionary changes in energy producing pathways, including fatty acid degradation and glycolysis/gluconeogenesis, in which fatty acids and glucose serve as the main energy resource, respectively. We found three Cetacea PSGs (*ACADVL, EHHADH*, and *HADHB*) in β-oxidation (GO: 0006635). These three PSGs encode four enzymes including acyl-CoA dehydrogenase, enoyl-CoA hydratase, 3-hydroxyacyl-CoA dehydrogenase, and thiolase which drives the β-oxidation (Bartlett and Eaton 2004). Our domain analysis revealed that cetacean specific amino acid substitutions occurred in the functional domains of all the four enzymes (Fig. 3, Table 2, and also Fig. S1, S9, and S13). Given the signatures of selection in the four main steps of this process, the β-oxidation of Cetacea may have been extensively modified during aquatic adaptation, potentially increasing the efficiency of producing energy from fatty acids. To our knowledge, this is the first report of an implication for evolution of Cetacea β-oxidation, including genomic and physiological studies.

The Cetacea PSGs exhibited three genes (*ACSS2, FBP1*, and *PDHB*) that were associated with the glycolysis/gluconeogenesis pathway (GO: 0006096 and GO: 0006094). Interestingly, gene expression of *FBP1* is regulated by bile acids (Yamagata et al. 2004), implying a possible functional connection with the other identified PSG, *CYP8B1. FBP1* encodes an enzyme at the rate limiting step of gluconeogenesis. Although metabolic studies of carnivores have primarily used domestic cats (Schermerhorn 2013), the selection of the Cetacea gluconeogenesis may provide a new insight into the mechanisms for glucose biosynthesis by carnivorous animals. The remaining two PSGs encodes enzymes involved in Acetyl-CoA synthesis: *PDHB* is a part of the pyruvate dehydrogenase complex that converts pyruvate into acetyl-CoA (Saunier et al. 2016), and *ACSS2* is a part of acetyl-CoA synthetase that catalyzes the formation of acetyl-CoA from acetate (Fujino et al. 2001). Acetyl-CoA is an important molecule that participates in the citrate cycle. It is noteworthy that we identified multiple signatures of positive selection involved in the synthesis of acetyl-CoA and selective pressure on β-oxidation. This emphasizes the importance of energy production for aquatic adaptation.

## Conclusion

The comparative analyses between aquatic and terrestrial Cetartiodactyla provided genetic signatures of the evolution of lipid metabolism in cetaceans, supporting divergent evolution of lipid metabolism since the divergence of these taxa from a common ancestor. This study is unique in that it used a sufficient number of both Cetacea and Bovidae samples in an equal manner that enabled the identification of molecular changes putatively reflecting their ecologies. The lineage-specific amino acid substitutions in domain regions implied functional modifications of multiple genes that are involved in important biological processes for aquatic adaptation. These genes and the associated pathways will be plausible targets for future investigations of cetacean lipid metabolism.

## Supporting Information

Additional Supporting Information may be found in the online version of this article at the publisher’s website:

## AUTHOR CONTRIBUTIONS

Y.E. conceived of the study, designed the study, and carried out the data collection, data analysis, bioinformatics works, and drafted the manuscript; K.K. and M.M. coordinated the study and participated in the design of the study, and helped draft the manuscript. All authors gave final approval for publication.

## ACKNOWLEDGEMENTS

The authors wish to thank Takushi Kishida, Daisuke Muramatsu, and Oliver A. Ryder for assisting with the data analysis and interpretation, and Maegan Fitzgerald for English correction.

This work was supported by the Japan Society for the Promotion of Science (JSPS; 16K14660 to K.K. and 25118005 to M.I-M.). This study was also partially supported by Kyoto University Supporting program for interaction-based initiative team studies (SPIRITS) to M.I-M. The WPI-iCeMS was supported by the World Premier International Research Centre Initiative (WPI), MEXT, Japan.

